# Sequence determinants of *in cell* condensate assembly morphology, dynamics, and oligomerization as measured by number and brightness analysis

**DOI:** 10.1101/2021.04.18.440340

**Authors:** Ryan J. Emenecker, Alex S. Holehouse, Lucia C. Strader

## Abstract

**Background:** Biomolecular condensates are non-stoichiometric assemblies that are characterized by their capacity to spatially concentrate biomolecules and play a key role in cellular organization. Proteins that drive the formation of biomolecular condensates frequently contain oligomerization domains and intrinsically disordered regions (IDRs), both of which can contribute multivalent interactions that drive higher-order assembly. Our understanding of the relative and temporal contribution of oligomerization domains and IDRs to the material properties of in vivo biomolecular condensates is limited. Similarly, the spatial and temporal dependence of protein oligomeric state inside condensates has been largely unexplored in vivo.

**Methods:** In this study, we combined quantitative microscopy with number and brightness analysis to investigate the aging, material properties, and protein oligomeric state of biomolecular condensates in vivo. Our work is focused on condensates formed by AUXIN RESPONSE FACTOR 19 (ARF19), which is a transcription factor integral to the signaling pathway for the plant hormone auxin. ARF19 contains a large central glutamine-rich IDR and a C-terminal Phox Bem1 (PB1) oligomerization domain and forms cytoplasmic condensates.

**Results:** Our results reveal that the IDR amino acid composition can influence the morphology and material properties of ARF19 condensates. In contrast the distribution of oligomeric species within condensates appears insensitive to the IDR composition. In addition, we identified a relationship between the abundance of higher- and lower-order oligomers within individual condensates and their apparent fluidity.

**Conclusions:** IDR amino acid composition affects condensate morphology and material properties. In ARF condensates, altering the amino acid composition of the IDR did not greatly affect the oligomeric state of proteins within the condensate.

## Background

Across all kingdoms of life, cells must accomplish the difficult task of organizing their intracellular environment. Cells can accomplish this through two primary mechanisms; compartmentalization through use of membrane-bound organelles or through the formation of biomolecular condensates [1–4]. Biomolecular condensates are non-stoichiometric assemblies of biomolecules that are defined by their common feature of spatially concentrating cellular components [3]. Growing evidence suggests that in many cases, biomolecular condensates form through the process of phase separation [4–8]. Of fundamental importance to this process is the multivalency of the molecules that undergo phase separation [8–10]. Multivalency refers to the capacity of a molecule to simultaneously engage in multiple intermolecular interactions. The ability of multivalent biomolecules to undergo coordinated and regulatable assembly is at the heart of biomolecular condensate formation, regardless of if the underlying mechanism is phase separation or some other process.

Many proteins that are capable of undergoing phase separation contain intrinsically disordered regions (IDRs) [7,11–14]. IDRs are protein regions that do not adopt a fixed three-dimensional structure but instead exist as an ensemble of conformations that interconvert between one another [15–17]. While IDRs are not strictly necessary for phase separation, in a number of specific biological systems IDRs have been found to be necessary and/or sufficient for phase separation and condensate formation [12,18–21]. However, folded domains often play key roles in facilitating initial oligomerization that licenses subsequent phase separation and can in their own right undergo phase separation absent IDRs [10,22–34]. Taken together, it should be clear that the molecular basis for multivalency is not constrained to a specific class of biomolecule.

The observation that IDRs can drive phase separation is often taken as evidence that they have evolved to facilitate biomolecular condensate formation. An alternative interpretation is that even if condensate formation is driven by folded domains, multivalent IDRs may be essential for the formation of dynamic, labile, and functionally responsive biomolecular assemblies [35,36]. Under this model, IDRs offer a means to encode locally tethered molecular lubricants that prevent aberrant assembly of folded domains and to tune the material state of biomolecular condensates [20,27,36–39]. With this in mind, understanding how structurally or chemically orthogonal multivalent interactions can tune condensate material properties represents an emerging set of questions.

Condensates are frequently well-described as viscoelastic materials, meaning they have an elastic response upon deformation and will also drip, flow, or wet like a viscous fluid [6]. The apparent viscosity of a condensate can range from liquids that rearrange in milliseconds to viscous solid-like assemblies that fuse on the order of hours or longer [40,41]. A growing body of literature supports an emerging view that condensate material state can be a key determinant of biological function [38–40,42–44]. With this in mind, a molecular understanding of interactions that determine material state represents an important next step in our ultimate goal of relating protein sequence and structure to cellular function. The viscosity of a condensate depends on the lifetime and the number of cohesive multivalent interactions that are responsible for assembly [7,9,45]. Previous work focused on IDRs has shown that amino acid composition and patterning can influence condensate dynamics in a manner that alters these parameters [20,46–48]. However, many phase separating proteins possess a modular protein architecture that include (at a minimum) an oligomerization domain and an IDR [10]. For these modular proteins, the relative impact of folded vs. disordered domains on the emergent material properties of a condensate is less well studied.

A final confounding factor when considering how IDRs tune the material properties of condensates emerges from the observation that IDRs are inherently sensitive to their solution environment [49–54]. Given the crowded and complex milieu of macromolecules, osmolytes, and ions *in vivo*, one may expect that the material properties for a condensate measured *in vitro* to be substantially different, an expectation supported by numerous studies. As such, to understand physiologically relevant determinants of condensate material state, ideally measurements of condensate dynamics and protein oligomeric state would be made in live cells.

Here we leverage the previously characterized modular transcription factor AUXIN RESPONSE FACTOR 19 (ARF19) from *Arabidopsis thaliana* as a model system to probe the determinants of condensate properties and protein oligomeric state in cells. Auxin is a plant hormone that is involved in essentially all plant growth and developmental processes [55]. ARF19 condensate formation has been proposed to be a mechanism by which signaling through the auxin signaling pathway is attenuated [22]. Notably, disruption of ARF19 condensate formation has a dramatic impact on the expression of auxin-responsive genes, implicating ARF19 condensates as global remodellers of auxin-dependent transcription [22]. ARF19 is composed of a DNA-binding domain followed by a large glutamine-rich IDR and a C-terminal Phox Bem1 (PB1) oligomerization domain. Importantly, *in vivo*, we have previously shown that the oligomerization domain and the IDR are essential for condensate formation [22]. As such, ARF19 is a convenient model system to examine the contribution of the IDR to material properties of condensates formed by proteins containing both IDRs and oligomerization domains.

## Methods

### Plant Growth

Seeds were surface sterilized [56] and then suspended in 0.1% agar. Suspended seeds were then kept at 4°C for 2 days for stratification. After stratification, seeds were plated on plant nutrient (PN) medium [57] + 0.5% (w/v) sucrose solidified with 0.6% agar. Seeds were then grown for 1 week at 22°C under continuous light before being transplanted to soil. Once transplanted to soil, seedlings were grown under long day (16 hours light : 8 hours of dark) conditions for 2-3 more weeks before being used for protoplast generation.

### Protoplast Isolation

For all protoplast transfections, an *arf19, arf7* double mutant in the Col-0 background was used to minimize the risk of native ARF19 or ARF7 interactions impacting condensate formation. Protoplasts were isolated from 3-4 week-old-plants via the tape-method as described in [58] with slight modifications. Briefly, after the upper epidermal surfaces of the leaves were peeled, peeled leaves were incubated in enzyme solution (1% cellulase, 0.25% macerozyme, 0.4 M mannitol, 20 mM KCl, 20 mM MES, 10 mM CaCl_2_, 0.1% BSA) in 6-well plates as opposed to Petri dishes. In addition, peeled leaves were incubated with the enzyme solution for 60-90 minutes. Lastly, leaves were shaken at 60 RPM during incubation with the enzyme solution.

### Protoplast Transfection

Protoplast transfection followed methods as described in [58] with slight modifications. After resuspension in the MMg solution, 150 μl of protoplasts were mixed with 10-12 μg of plasmid DNA. Next, protoplasts were mixed with an equal volume of the PEG solution and rocked back and forth gently to mix the protoplasts with the PEG solution. Once mixed, protoplasts were incubated for 10-12 minutes before 660 μl of W5 solution was added. After protoplast transfection, protoplasts were immediately suspended in 2 ml of W1 buffer [58] and then dispensed onto 50 x 7 mm round bottom glass dishes (Ted Pella Inc, product number 14035-120). All expressions in protoplasts utilized the UBQ10 promoter, and ARF19 as well as the two ARF19 variants contained an N-terminal mVenus tag.

### Vector Construction

The transient protoplast expression vectors for ARF19 and ARF19 QtoS were made through recombination from a pENTR vector into pUBQ10:mVenus-GW and have been described previously [22]. For the ARF19 QtoG variant, the ARF19 QtoG IDR was synthesized with 8 base pairs of overhangs to the region immediately upstream of the ARF19 IDR at the 5′ end and 20 base pairs of overhang to the region immediately downstream of the ARF19 IDR at the 3′ end as a gBlock by Integrated DNA Technologies. Then a pENTR-ARF19 vector was linearized via PCR amplification using Pfx Platinum polymerase (Life Technologies) such that it no longer contained the wild-type IDR, and a 12 base pair overhang with the 5′ end of the QtoG IDR gBlock was added at this step using the primers 5′-AACTAGACTTAAACCAGGGAACATC-3′ and 5′-AACTTGGTTCCCAACTATGGC-3′. The resultant PCR product and the QtoG IDR gBlock were used in an In-Fusion cloning reaction (Takara) to generate pENTR-ARF19 QtoG IDR. After sequence confirmation, pENTR-ARF19 QtoG IDR was recombined into the pUBQ10:*mVenus-GW* vector using LR Clonase II (ThermoFisher) to generate pUBQ10:mVenus-ARF19 QtoG, which was subsequently sequence confirmed.

### Microscopy Imaging

Immediately after transfection, protoplasts suspended in 2 ml of W1 were dispensed onto Ted Pella Inc 50×7 mm PELCO Round Bottom Dishes (glass, 40 mm) (product number 14035-120). The protoplasts were then incubated for approximately 16 hours in the round bottom dish with a vacuum grease sealed lid enclosing the dish such that the dish did not dry out and alter the concentrations of solutes prior imaging. After approximately 16 hours, the lid was removed from the dish and the protoplasts were placed on the confocal stage for imaging. Imaging was carried out using the Leica SP8 confocal microscope. All images of condensates used the HyD detector and a 40x water immersion lens. All images in Figure 1C used the Leica Lightning Imaging module and Lightning deconvolution. For the time lapses of condensate fusion events in figure 4A, images were obtained from time lapses of individual whole protoplasts. Unlike the images presented in Figure 1C, the images in figure 4A did not use the Leica Lightning module and used the 20x, dry-immersion objective.

**Figure 1.**
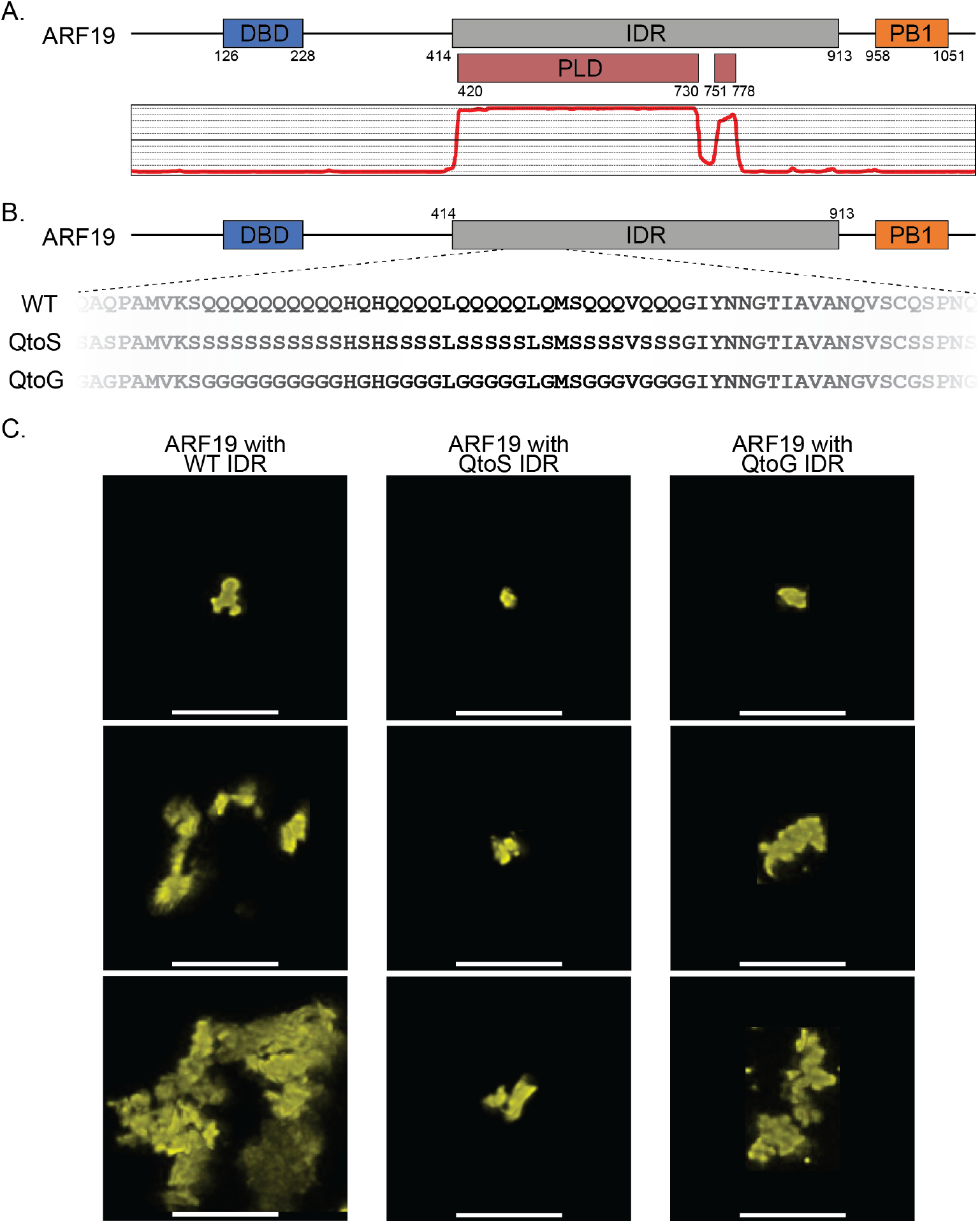
Altering composition of the ARF19 IDR impacts the morphology of ARF19 condensates. **(A)** Schematic of the ARF19 protein showing the location of the DBD and PB1 domains. The graph below shows predicted prion-like domains within the ARF19 IDR, which were predicted using PLAAC [61]. **(B)** Schematic showing a subsection of the ARF19 IDR highlighting differences in IDR composition for the QtoS (middle) or QtoG (bottom) variants. While this schematic only shows a subsection of the IDR, for the QtoG or QtoS variants, all glutamines were changed to glycine or serine, respectively. **(C)** Images showing the range of condensate morphologies formed by wild-type ARF19 or ARF19 with the altered IDR compositions. Images were chosen to represent the breadth of condensate morphology and sizes that are frequently observed across protoplasts when expressing the various constructs. All images were taken in different protoplasts approximately 16 hours after transfection. Scale bars represent 5 microns.

### Number and Brightness Imaging and Analysis

All imaging for N&B was taken on a Leica SP8 using a HyD detector in photon counting mode using a 40x water immersion objective. Prior to imaging, slight adjustments to the correction collar on the objective were made as needed to minimize differences in acquired signal due to differences in glass thickness of the round bottom dishes. The laser used was a 514 nm set at 0.01% power, and the range of wavelengths used for capturing the images were those between 519 nm and 550 nm. The pinhole was set at 1 AU, the scan speed was 310 Hz, and the zoom was set at 11.5. One hundred consecutive frames were captured for each data point, and the interval between each frame was 0.839 seconds (1 minute, 23.871 seconds total acquisition time). Pixel dwell time was 8.19 μsec. Images were captured at a 256 x 256 format. For analysis, we used the SimFCS software [59]. We used protoplasts expressing mVenus under the UBQ10 promoter in order to calibrate software parameters such as the S-factor. Specifically, the region that quantified monomer brightness was determined using protoplasts expressing mVenus by selecting the fluorescence-positive pixels with a cursor. Using the assumption that anything twice the brightness of the calibrated monomer would be a dimer, three times the brightness would be a trimer, and so forth, the amount of higher order oligomers was quantified. For our analyses, monomers, dimers, 3-10mers, and 10+-mers were quantified with the exception of Figure 5A. For Figure 5A, analysis of the different oligomeric species in the condensates was carried out similar to as before with the exception that individual species from monomers to decamers and 10+-mers were individually analyzed in order to obtain images showing their distribution within each condensate. Then, the images were uploaded to Adobe Illustrator and overlaid with one another. Finally, each pixel was false colored such that each pixel corresponded to the oligomeric species identified in the N&B analysis.

### FRAP Imaging and Analysis

FRAP imaging was carried out on a Leica SP8 using a PMT detector and a 40x water immersion objective using the Leica FRAP module. All FRAP imaging was carried out immediately after N&B imaging. Pre- and post-photobleaching image acquisition used a 514 nm laser at 0.06% power with a range of acquisition of 519 nm to 550 nm. All imaging used a 512×512 format and a scan speed of 1400 hz. One pre-bleach image was acquired followed by the photobleaching and then 120 post-bleach images were captured at 1 second intervals. The duration per acquisition of each image was 0.371 seconds. For FRAP imaging, the zoom was adjusted as needed depending on the size of the condensate. All optional SP8-specific FRAP module settings were set as follows: fly mode -off, zoom in - on, change bleach format - off, set background to zero - off, delete bleach images after scan - off. For photobleaching, the 448 nm, 488 nm, 514 nm, and 552 nm lasers were set to 100% power and targeted to approximately one half of the condensate for a total of 1.8762 seconds. Following image acquisition, data was imported into FIJI (FIJI Is Just ImageJ) [60] in the original .lif file format for analysis. The percent recovery was determined by quantifying the amount of recovery observed in the photobleached region post-photobleaching.

## Results

### Altering IDR composition dramatically impacts intracellular condensate morphology and dynamics

ARF19 contains a ∼500-residue glutamine-rich IDR that includes a large prion-like domain (PLD), a class of low-complexity IDR enriched in polar amino acids [61,62]. The ARF19 IDR lies between an N-terminal DNA binding domain and a C-terminal PB1 oligomerization domain (Figure 1A). Given its large size and strong sequence bias, we wondered if changing the amino acid composition (while maintaining the natural enrichment for polar amino acids) would alter condensate properties.

To explore how IDR composition impacts condensates formed by ARF19 *in vivo*, we took advantage of a transient expression protoplast system. Protoplasts are individual spherical cells in which the cell wall has been removed through enzymatic degradation. For our studies, we used protoplasts derived from leaf mesophyll cells isolated from three-week-old *A. thaliana*. Importantly, using protoplasts allowed us to examine the behavior of the ARF19 condensates within the cellular environment. In addition to offering an *in vivo* environment, protoplasts provide a convenient system to examine condensates for several reasons. The large size of protoplasts makes it easy to image the condensates, their ease of transfection makes it easy to examine many different condensate-forming proteins, and the ability to detect protein expression after transfection in as little as 90 minutes allows one to quickly examine the events leading up to condensate formation. In addition, due to the tight temporal control that transient expression affords, it is possible to estimate condensate age. This enables us to examine condensate formation at a single time point after transfection, an important feature given some condensate properties can change dramatically in a time-dependent manner.

We examined the morphology of condensates in protoplasts formed by full-length ARF19 with a wild-type IDR, an IDR where all glutamines were changed to glycines (QtoG) or an IDR where all glutamines were changed to serines (QtoS) (Figure 1B). Glycine and serine were chosen here as amino acids frequently found in condensate-forming IDRs that retain the polar chemistry of glutamine sidechains yet alter steric and physicochemical properties [21]. Condensates formed by our three constructs displayed striking differences in morphology (Figure 1C). Condensates formed by the WT IDR were in general large, amorphous multilobed assemblies in line with our previous work [22]. In contrast, QtoS condensates were smaller and more spherical, whereas QtoG condensates were intermediate in terms of morphology. Given the oligomerization domain is necessary for condensate formation in plants when expressed at physiological levels, these results reveal that condensate formation and condensate morphology can be uncoupled from one another [22].

Condensate morphology is inherently linked to condensate dynamics. For condensates with liquid-like properties, condensate morphology favors spherical assemblies that minimize the interface between the dense and dilute phases. In contrast, for solid-like condensates, morphology is dictated by intramolecular interactions that are inherently anisotropic, giving rise to amorphous assemblies, networked solids, sheets, cross-linked polymers, or any number of non-spherical assemblies [14,25,39,63–66]. However, spherical condensates that rapidly mature from liquid to solids have been observed in many systems, with a ‘bunch of grapes’ type architecture typifying systems in which arrested spherical assemblies adsorb onto one another [27,67]. As such, we suspected that the dynamics of the three variants were likely impacted by the differing IDR compositions. For example, the more spherical morphology of the QtoS condensates may imply enhanced dynamics compared to the more irregular WT condensates.

To assess this, we examined the dynamics of the different ARF19 variants using half-condensate Fluorescence Recovery After Photobleaching (FRAP). Importantly, because in many cases condensate undergo a time-dependent loss in dynamics [41,44], all FRAP measurements were carried out approximately sixteen hours after protoplast transfection.

Condensates formed from all three IDR variants showed relatively low levels of fluorescence recovery. Consistent with prior *in planta*, wild-type ARF19 condensates exhibited minimal recovery after photobleaching in protoplasts with an average percentage recovery of just ∼7% after two minutes (Figure 2). In contrast, we found that the QtoS and QtoG IDR variants exhibited slightly more liquid-like properties, both having ∼9.6% recovery after two minutes. Despite previous examples where changing glutamine to glycine resulted in more liquid-like condensate dynamics, we found that the QtoG variant was only slightly more liquid-like than wild-type (Figure 2A, 2B) [20]. Nonetheless, this demonstrates that the IDR can influence the morphology and dynamics of ARF19 condensates despite the essential role of the oligomerization domain in their assembly.

**Figure 2.**
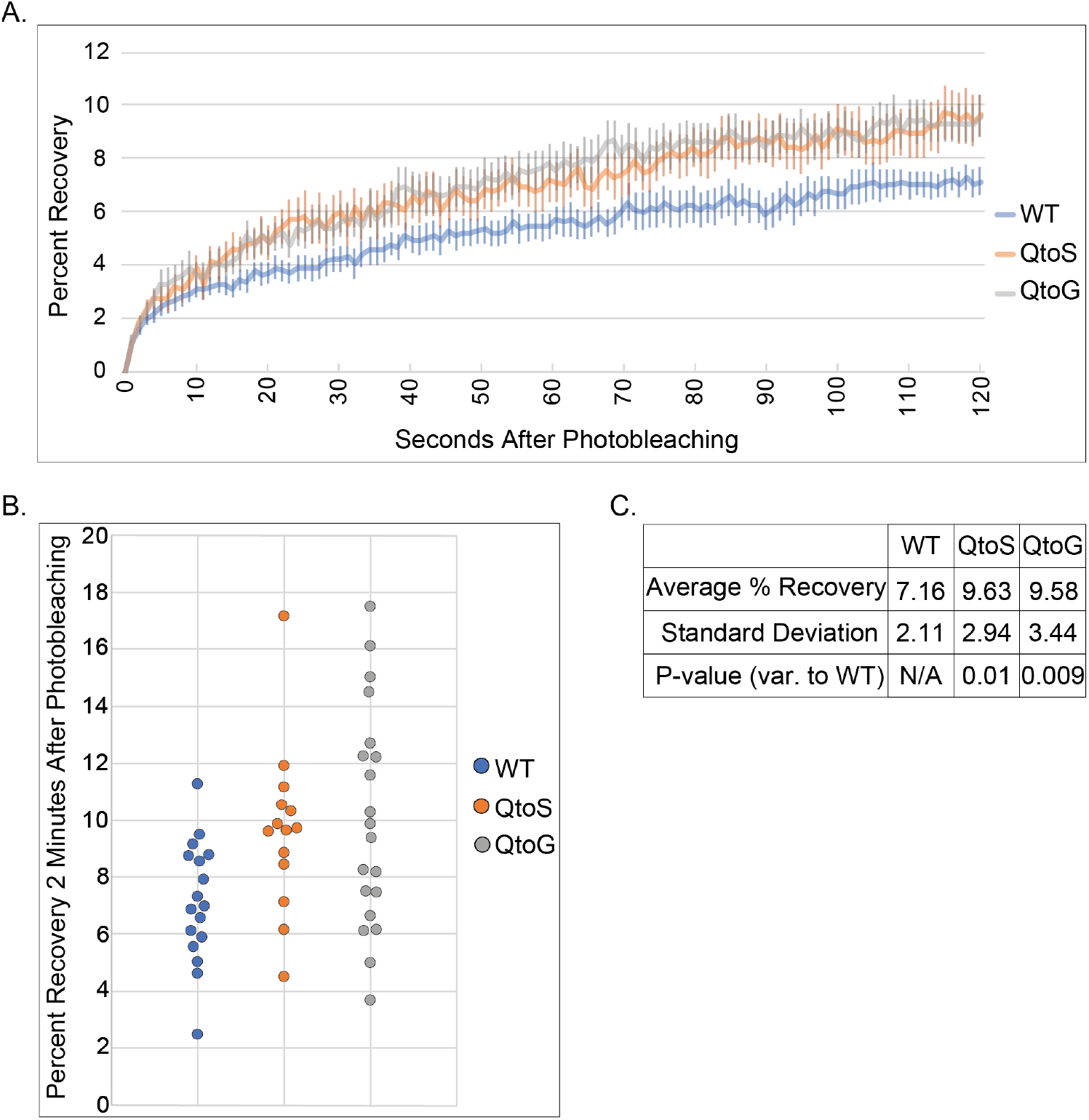
IDR composition impacts the fluidity of ARF19 condensates. **(A)** Average fluorescence recovery curves for the three ARF19 variants. Error bars report standard error of the mean. **(B)** Each point represents the percent recovery 2 minutes post-photobleaching of an individual condensate. For all FRAP experiments, only one half of the condensate was photobleached. N=17 (WT), N=14 (QtoS), N=20 (QtoG). **(C)** A table summarizing data from panel A. P-values were calculated using a two-sided t-test comparing the values from wild-type to each of the two IDR variants.

### Number and Brightness analysis reveals oligomeric state of ARF19 condensates

As a means of interrogating the impact that IDR composition has on the oligomeric state of the ARF19 condensates, we utilized number and brightness (N&B) analysis. N&B is a fluorescence microscopy method that uses a series of images taken over time to measure the average number of molecules and their oligomeric state in each pixel [68]. In this context, the term oligomeric state describes whether a given molecule behaves as a single unit (i.e. a monomer) or as a higher order assembly (i.e., dimers, trimers *etc*.).

As with the FRAP measurements, the N&B measurements were taken approximately sixteen hours after protoplast transfection. For each condensate, we quantified the percentage of monomers and dimers (termed ‘lower order’ oligomers) and the percentage of oligomers equal to ten or higher (termed ‘higher order’ oligomers).

To our surprise, regardless of whether the percentage of monomers and dimers or the percentage of 10+-mers were examined, there was very little difference in the average oligomeric state between the three variants (Figure 3A, 3B, 3C). This is in contrast to our FRAP measurements where we saw an increase in the average fluorescence recovery of condensates formed by the QtoS and QtoG variants. These results support a model whereby IDR composition has the capacity to influence the morphology and dynamics of the ARF19 condensates but not the oligomeric state.

**Figure 3.**
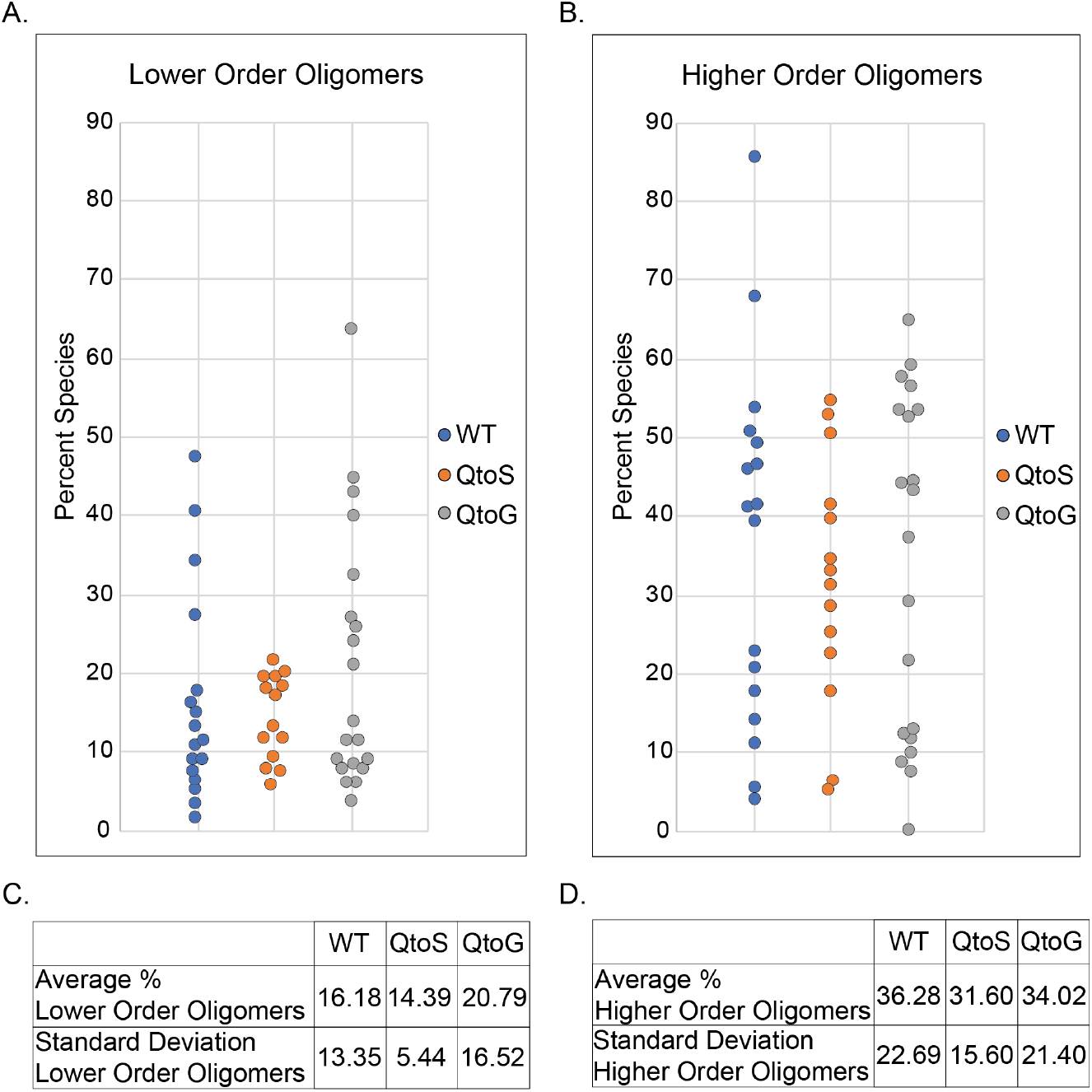
IDR composition has little impact on the oligomeric state of ARF19 condensates. **(A)** Each point represents the percent of monomers and dimers out of the total number of measured oligomers for an individual condensate. **(B)** Each point represents the percent of 10+-mers out of the total number of measured oligomers for an individual condensate. **(C)** A table summarizing the data from figure panel A. **(D)** A table summarizing the data from figure panel B. Statistical testing is not shown because none of the comparisons were statistically significant.

Biomolecular condensates frequently undergo a time-dependent maturation in their material state. Condensates often form with liquid-like properties but may undergo a persistent loss of dynamics over time in a process referred to as aging, maturation, or as a liquid-to-solid transition [41,44,69,70]. In investigating the relationship between oligomeric state and condensate dynamics in ARF19, we found that the ARF19 condensates also exhibit a decrease in apparent fluidity over time. Time lapses of protoplasts immediately after transfection show that the condensates initially readily fuse with one another suggesting they are liquid-like in nature (Figure 4A). In contrast, over time the condensates become more solid and are unable to fully fuse resulting in the formation of condensates with apparent substructure (Figure 4A). With this observation, we sought to determine if the oligomeric state of ARF19 inside condensates showed an analogous time-dependent maturation. Unlike condensates measured after sixteen hours, we found that newly formed ARF19 condensates contain few, if any, higher-order oligomers (Figure 4B, C). This suggests that there may be a relationship between the apparent fluidity and the accumulation of higher order oligomers within individual condensates.

**Figure 4.**
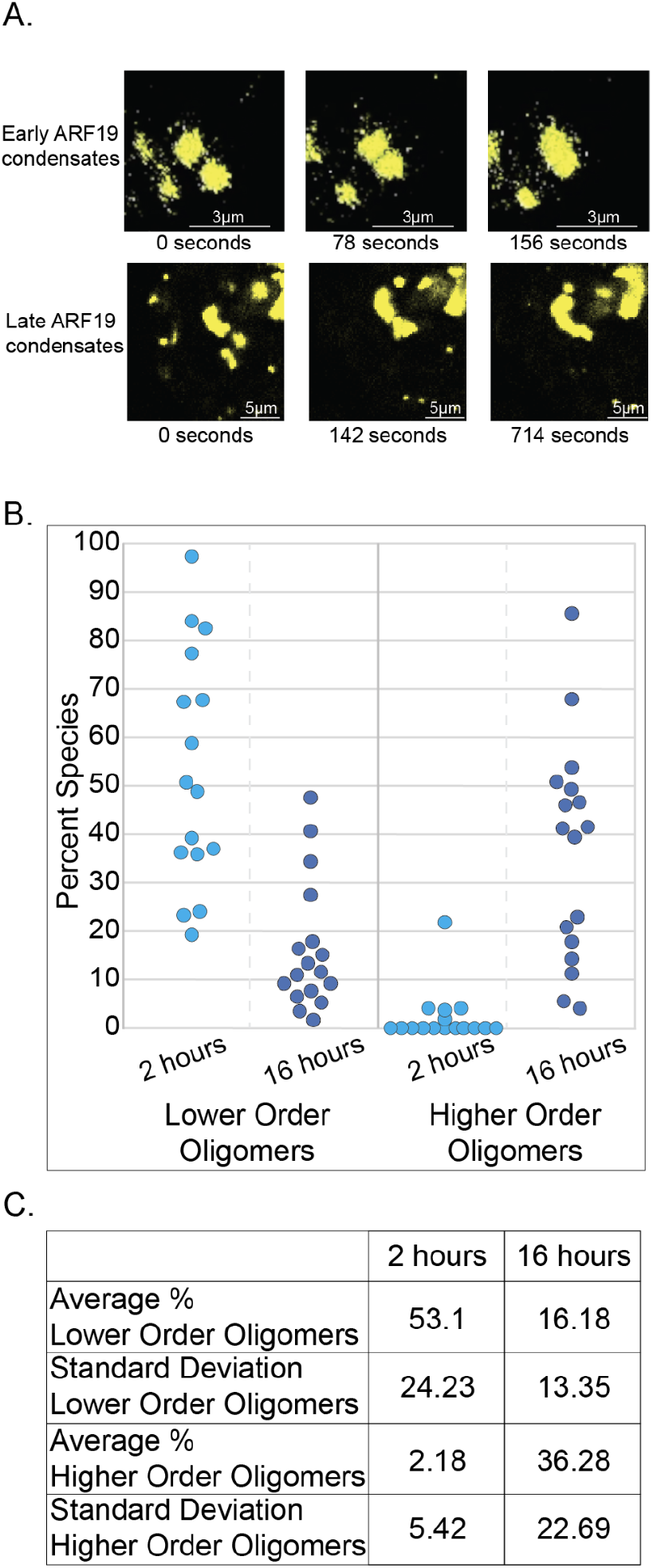
ARF19 condensates are initially liquid-like and lack higher-order oligomers. **(A)** Time lapse images of wild-type ARF19 condensates in protoplasts. The top images are from the beginning of a time-lapse series started shortly after protoplast transfection (early ARF19 condensates) and show an example where the condensates have liquid-like behavior. The bottom images are from a later time point in a time lapse series (late ARF19 condensates) and show an example where the condensates appear to partially fuse but ultimately are unable to fully fuse resulting in a ‘grape-bunch’ like morphology. Note, time intervals between the bottom panels are not equal from panel to panel. (**B)** Each point shows the percent values for either lower order oligomers (left) or higher order oligomers (right) for individual ARF19 condensates approximately two or sixteen hours after protoplast transfection N=16 (3 hours) and N=17 (16 hours). **(C)** Table summarizing the data from figure panel B.

Apart from imparting information with respect to the quantity of different oligomeric species within condensates, N&B analysis can also reveal information with respect to the spatial distribution of oligomers within a condensate. N&B analysis has previously revealed the distribution of oligomers within ARF19 condensates, finding that the higher order oligomers tend to be towards the center of the condensates and the monomers and dimers can be found towards the periphery [22]. After we found that the abundance of lower order and higher order oligomers is dramatically different between the early and late ARF19 condensates, we sought to examine whether the spatial distribution of lower and higher order oligomers was different between the early and late condensates. We found that the distribution of oligomers in early condensates was similar to that seen in the later condensates (Figure 5A). Therefore, while the relative abundance of different oligomeric species differs significantly between the early and late ARF19 condensates (Figure 5B), the general pattern where higher order oligomers tend to exist towards the center of the condensates and the lower order oligomers towards the periphery appears to be consistent over time.

**Figure 5.**
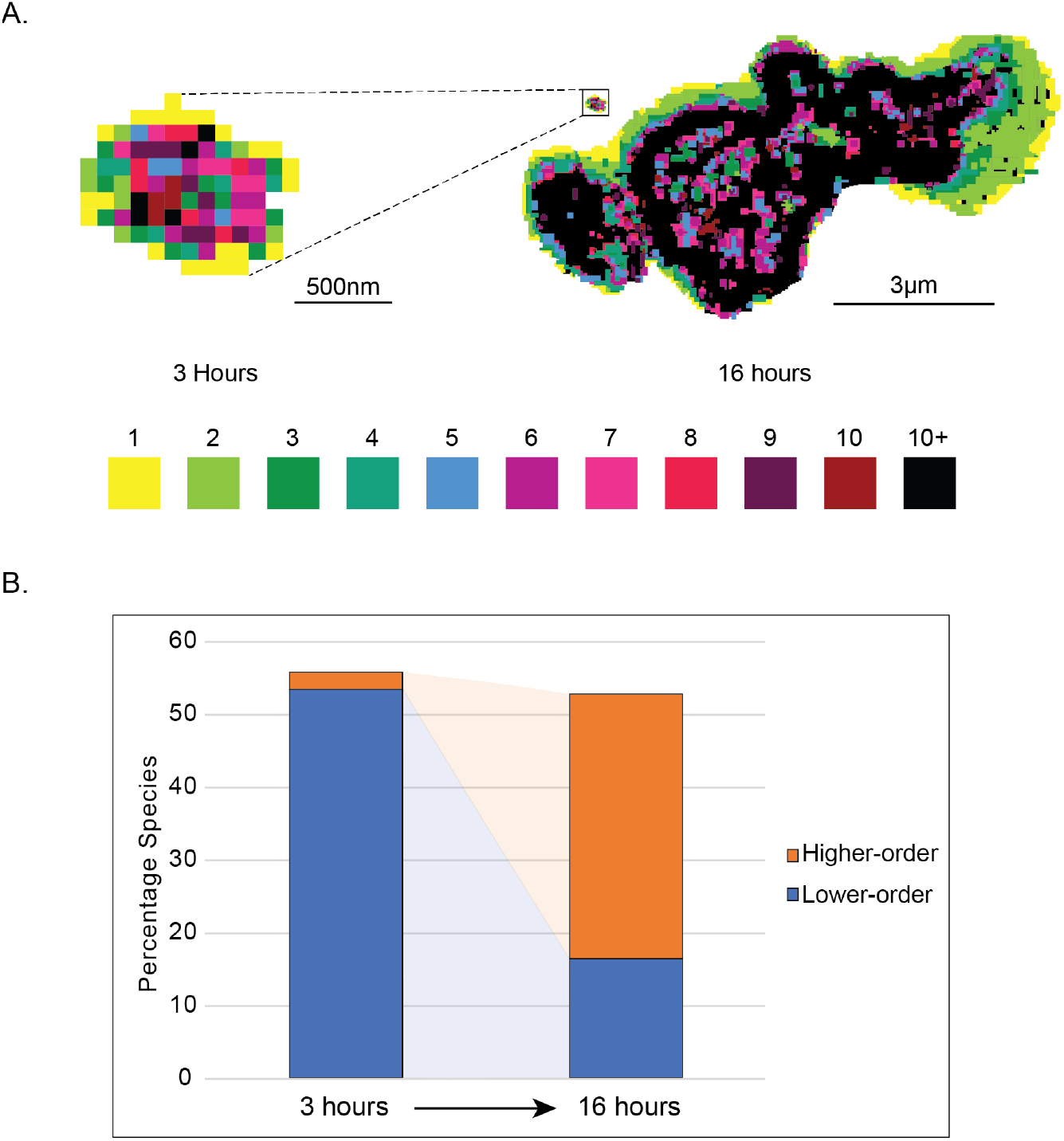
Distribution of oligomeric species in ARF19 condensates over space and time. **(A)** N&B analysis showing the spatial distribution of various oligomeric species in early (left) and late (right) ARF19 condensates. Each oligomeric species corresponds to a different color. The size of the early condensate relative to the late condensate can be seen in the box towards the top left of the late condensate. **(B)** N&B analysis showing the average percentage of higher-order and lower-order oligomers in ARF19 condensates in early condensates (left) and late condensates (right).

### Examining the relationship between oligomeric state and fluorescence recovery of individual condensates

Finally, we examined the relationship between condensate fluidity and oligomeric state. During data acquisition we carried out N&B followed by FRAP measurements on the same individual condensates, allowing us to directly correlate oligomeric state with condensate dynamics on a per-condensate basis. We found that the condensates formed by wild-type ARF19 showed a weak but positive correlation between the percent of lower order oligomers or of higher order oligomers and the percent recovery of individual condensates (Figure 6A, B). In contrast, the QtoS and QtoG variants showed very little correlation between either the percent of lower or higher order oligomers and fluorescence recovery for individual condensates. (Figures 6C-F).

**Figure 6.**
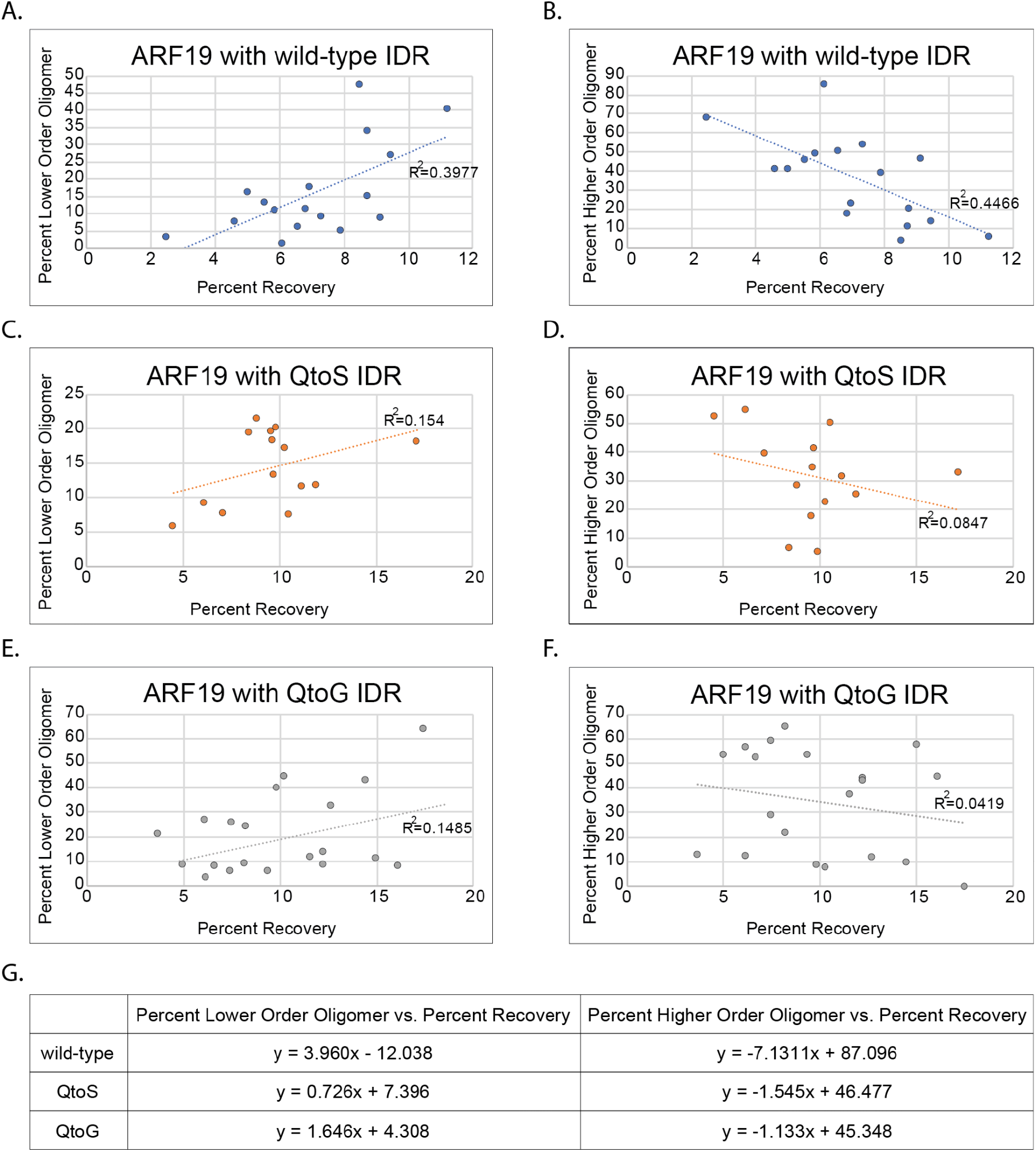
IDR composition-dependent relationship between condensate oligomeric state and fluidity. **(A-F)** Each point shows N&B and FRAP data for an individual condensate. Panels on the left show the relationship between the percent of lower order oligomers and the fluorescence recovery two minutes post-photobleaching for individual condensates whereas panels on the right show the relationship between higher order oligomers and percent recovery for individual condensates. Dashed lines are linear fit lines. R^2^ values are shown near the linear fit lines. N=17 (WT), N=14 (QtoS), and N=20 (QtoG). **(G)** Equations for linear fit lines shown for each graph.

## Discussion

Biomolecular condensates have emerged as key organizers of cellular matter, whereby multivalent interactions underlie the assembly, recruitment, and regulation of a wide array of cellular bodies. The perspective that condensate formation is driven by intrinsically disordered regions has given way to a broader appreciation for distributed multivalency, whereby both disordered regions and folded oligomerization domains can make key contributions to the assembly of condensates [9]. Furthermore, condensate material state is increasingly being recognized as an important contributor to cellular function, either directly or indirectly [38–40].

In this study, we utilized ARF19 as a system to examine the role that IDR composition has on the emergent material properties of ARF19 condensates. Importantly, because of the inherent sensitivity that IDRs and more generally phase separation have to the surrounding solution environment, we carried out this work inside of protoplasts, which are plant cells that lack an external cell wall [11,21]. This allowed us to examine the effect that IDR composition has on condensate material properties within the cellular environment. In the context of material properties, this is of particular importance as there are known examples where condensates that exhibit liquid-like dynamics *in vivo* form non-dynamic, solid assemblies *in vitro*, suggesting that the solution environment has a direct influence on the material properties of condensates [20,44]. By first quantifying oligomeric state using Number and Brightness (N&B) analysis and then examining fluidity using Fluorescence Recovery After Photobleaching (FRAP) for ARF19 variants with differing IDR compositions, we were able to assess the relationship between IDR composition and the apparent solidity and oligomeric state of ARF19 condensates. In addition, because we carried out these analyses sequentially for individual condensates, we were able to examine the relationship between the oligomeric state of the condensates and condensate fluidity, in an effective “single-condensate spectroscopy” type experiment.

Whereas the oligomerization domain of ARF19 promotes condensate formation through a well-defined binding interface, the mechanism by which the glutamine-rich IDR of ARF19 contributes is less obvious [22,71]. Extant work on polyglutamine has demonstrated its robust tendency to undergo self assembly, and previous studies have shown that the glutamine content in an IDR can impact the propensity of condensates to undergo time-dependent maturation and loss of dynamics [20,72–76]. In addition, prion-like domains, which are generally associated with aggregate or condensate formation, are frequently glutamine-rich [61,62,77–79]. In regards to the underlying mechanism by which glutamine contributes to material properties of condensates, glutamine-rich IDRs can form coiled-coils that have the capacity to facilitate protein-protein interactions and multimerization [76,79,80]. Therefore, in the context of ARF19, which has a glutamine-rich IDR with multiple polyglutamine stretches, the glutamine content may underlie the material properties of ARF19 condensates.

IDRs are frequently found to be both necessary and sufficient for biomolecular condensate formation. Despite their clear importance in condensate formation, little is known about the impact that IDR composition has on the emergent material properties of condensates *in vivo*. In addition, while multiple examples of proteins requiring both their IDR and oligomerization domain to form condensates have emerged, how IDR composition impacts condensate dynamics in this context is poorly understood. Here, through use of FRAP and Number and Brightness analysis (N&B) coupled with altering the composition of the IDR in ARF19, we have begun to shed light on this question.

### IDR composition influences condensate dynamics

From our FRAP analysis of condensates formed from wild-type ARF19 or our ARF19 variants, we found that the QtoS and QtoG variants resulted in condensates with slightly increased fluidity. While the IDR can tune condensate properties, it is worth noting that in all three cases ARF19 condensates were relatively solid at 16 hours post-transfection, reaching a maximum average fluorescence recovery after two minutes of ∼9.6% (QtoS and QtoG variants, Figure 2).

The behavior of the QtoG variant is in contrast to a previous study which found that changing glutamine to glycine in the FUS PLD resulted in the formation of more dynamic condensates that did not undergo time-dependent maturation [20]. There are numerous possible explanations for this discrepancy. First, the previous study assessed condensate fluidity *in vitro* whereas our QtoG variant was examined within the cellular environment. Secondly, FUS and ARF19 differ in more than just their IDRs, and multivalent interactions facilitated by other protein regions that differ between the two proteins may also contribute to the differing results. Finally, the composition and patterning of amino acids in the FUS PLD and the ARF19 PLD are substantially different, with the FUS PLD containing an abundance of tyrosine residues that are absent in the ARF19 PLD. As such, the differences between the two outcomes may simply be due to the different IDR compositions. Moreover, while both are glutamine rich, unlike the ARF19 PLD, the FUS PLD lacks contiguous glutamine tracts. Prior work has established that polyglycine shows poor solubility in water and exists in compact, collapsed conformations that can undergo self-assembly [81–84]. As such, the acquisition of polyglycine tracts in the QtoG variant may be an additional determinant that contributes to differences in assembly behavior compared to the FUS PLD variant.

Glutamine rich sequences have been shown to form coiled-coiled domains that drive oligomerization through a conditionally-structured interface [80,85,86]. Moreover, analysis of the ARF19 sequence with the COILS webserver predicts a coiled-coil domain in line with one of the glutamine-rich subregions in the IDR (Figure S1) [87]. However, we can largely exclude coiled-coils as a key determinant of protein oligomeric state given that glycine strongly impedes helix formation [88,89]. With this in mind, the QtoG variant should fundamentally prevent any possible coiled-coil association, yet no different in intra-condensate oligomeric state is observed across our three variants (Figure 3). While we cannot rule out the possibility that coiled-coil domains may influence condensate morphology, it is conceptually challenging to envisage a model in which the presence or absence of a coiled-coil domain does not influence oligomeric state yet alters higher-order assembly. As such, we interpret our results to mean the glutamine-rich IDR functions in a largely unstructured manner.

The increased recovery dynamics observed in the QtoS variant may reflect various possible molecular origins. This result could report on a reduction in IDR-mediated interactions due to the replacement of a secondary amide sidechain (glutamine) for a hydroxyl group (serine). Alternatively it may reflect a change in residual structure, as implicated by work that suggests poly-serine might adopt a more expanded, rigid conformation [90]. Nevertheless, more work is needed to extrapolate results from simple homopolymeric peptides to observations in the context of full-length proteins.

ARF19 is part of a family of 23 ARFs in Arabidopsis that is broken into three separate clades [55]. The only ARFs currently known to form condensates in plants belong to clade A. These ARFs have characteristic glutamine-rich IDRs and are thought to act as transcriptional activators [91,92]. In contrast, there is currently no evidence that clade B ARFs form condensates in plants. Clade B ARFs are thought to be transcriptional repressors and contain serine-rich IDRs [91,92]. Our initial decision to alter the ARF19 IDR to become serine rich was based on the observation that none of the clade B ARFs, which contain serine-rich IDRs, are known to form condensates in plants. However, the minimal differences in solidity seen in our QtoS variant when compared to wild type ARF19 may suggest that the serine-rich nature of the clade B ARFs is not the inherent reason why these ARFs do not form condensates in plants.

### Oligomeric state is insensitive to IDR composition

In contrast to our FRAP data, we were unable to identify significant differences in the oligomeric state between any of the ARF19 variants. We interpret this to mean that while the IDR has the capacity to modulate the material properties of ARF19 condensates, the oligomeric state may be less impacted or all together independent of IDR composition. However, we cannot unambiguously conclude this due to the limited number of variants we analyzed, and the resulting limited statistical power.

Given that *in planta* both the IDR and the PB1 oligomerization domain are necessary for condensate formation, it is almost certain that the IDR contributes to the multivalent interactions that are essential for condensate formation. In support of this notion, when we expressed the PB1 domain of ARF19 alone in protoplasts, even among the protoplasts with the highest apparent accumulation of the protein, we did not observe any condensates, nor did we detect higher order oligomers through our N&B analysis (Table S1). In contrast, the PB1 domain forms multimers in vitro even in the absence of the IDR [71]. As such, our results support a model in which IDRs enhance the driving force for lower-order oligomers *in vivo*, but that this effect is sufficiently subtle that the QtoS and QtoG variants do not significantly perturb the effect *vis-à-vis* wild type. These results do not exclude the possibility that IDRs also stabilize higher order oligomers, a behavior we would expect to hold true. Future studies examining the oligomeric state of protein variants where IDRs are more dramatically altered, ideally in a way that minimizes the likelihood of the IDR contributing to multivalent interactions, should shed light on this question.

### Oligomeric populations change during condensate maturation in living cells

Given the observation that many condensates undergo time-dependent changes in dynamics and organization, we applied N&B analysis to assess how the oligomeric state of molecules inside condensates changes as a function of time. While condensates measured immediately after assembly were composed predominantly of monomers, after 16 hours we found a much larger population of higher order oligomers. These results are consistent with a model in which the high local concentration of molecules within a condensate drives concentration-dependent higher-order assembly, which in turn offers a structural explanation for changes in condensate dynamics. As larger oligomers form, their internal re-arrangement will become increasingly constrained due to molecular entanglement such that a jamming transition may ultimately occur, leading to a kinetically arrested assembly.

While our results here are readily interpretable in the context of oligomerization driven by the PB1 domain, the same principle is applicable to other systems in which distinct modes of assembly, such as the assembly of liquid-like condensates driven by distributed aromatic motifs, followed by a liquid-to-solid transition driven by the acquisition of structured cross-beta interactions [46,93–97]. Our previous work demonstrated that oligomeric state in condensates varies as a function of spatial position across the condensate, with more lower-order species on the surface and higher-order species in the interior [22]. Taken together, N&B analysis reveals that, at least for ARF19, oligomeric state varies in both space and time, revealing a rich and perhaps surprisingly complex oligomeric landscape of intra-condensate molecules.

### Oligomeric state can influence condensate dynamics

By carrying out N&B analysis followed by FRAP on individual condensates, we were able to establish a weak but clear correlation between the oligomeric state and the fluidity of condensates. This relationship is consistent with our finding that liquid-like ARF19 condensates examined shortly after formation do not contain substantial accumulations of higher order oligomers.

In contrast to wild-type ARF19 condensates, we did not observe a strong correlation between oligomeric state and condensate dynamics for condensates formed by either the QtoS or QtoG ARF19 variants. While it is possible that altering the IDR composition disrupted this relationship in some non-obvious way, it is also possible that the N&B data simply had too much noise for the QtoS and QtoG variants for us to see a clear relationship in this instance. Studying the behavior of biomolecular condensates *in vivo* is extremely challenging due to the inherent dynamic nature of the intracellular environment. In anecdotal support of this, in our protoplast system the ARF19 (or ARF19 variant) condensates were frequently highly mobile making capturing high-quality microscopy data challenging. Approximately 95% of the data acquired for this study had to be discarded prior to analysis simply due to the condensate moving out of view during the ∼4 minutes of data acquisition. Furthermore, given the substantially reduced size of condensates formed by the QtoS and QtoG variants, the likelihood of the variant condensates moving out of the Z-plane during acquisition was much higher than for wild-type condensates. This is not to say that *in vivo* studies of this type should not be attempted; rather, results such as the discrepancy between the presence of a relationship between the oligomeric state and the apparent fluidity of condensates formed by wild-type ARF19 and the lack of such a relationship in condensates formed by the QtoS and QtoG should be carefully considered. However, the prospect of ever-improving technologies that allow for more rapid and accurate acquisition of *in vivo* data will inevitably allow for a more accurate assessment of this relationship.

## Conclusions

In all, our work here offers direct insight into the relationships between IDR composition, condensate dynamics, and oligomeric state for a condensate-forming protein containing both an IDR and an oligomerization domain. Our results support an emerging consensus in which IDR composition impacts the emergent physical properties of biomolecular condensates both *in vitro* and *in vivo*. In contrast, our work suggests that, at least in the ARF19 system, IDR composition has a more limited role in governing the oligomeric state of *in vivo* condensates. Taken together, these results support a general model in which structurally and chemically orthogonal multivalent interactions can contribute distinct attributes to the emergent properties of biomolecular condensates.

## Author’s contributions

R.J.E., A.S.H., and L.C.S. designed this study and interpreted data. R.J.E. performed all described experiments. All authors read and approved the final manuscript.

## Acknowledgements

We thank Ishan Taneja for helpful comments on this manuscript.

