## Supplementary information for "Sequence determinants of *in cell* condensate assembly morphology, dynamics, and oligomerization as measured by number and brightness analysis"

Sequences of ARF19 and ARF19 variants. For all three variants, the region that was considered the predicted IDR (which is where glutamines were changed to G or S in the QtoG or QtoS variants, respectively) is underlined. The region in ARf19 wildtype that is in bold is the predicted prion-like domain **(PLD)**. The DNA-binding domain as annotated by Uniprot is in blue text. The phox/Bemp1 (PB1) domain as annotated by Uniprot is in orange text.

**>ARF19 wildtype**

MKAPSNGFLPSSNEGEKKPINSQLWHACAGPLVSLPPVGSLVVYFPQGHSEQVAASMQKQ
TDFIPNYPNLPSKLICLLHSVTLHADTETDEVYAQMTLQPVNKYDREALLASDMGLKLNR
QPTEFFCKTLTASDTSTHGGFSVPRRAAEKIFPPLDFSMQPPAQEIVAKDLHDTTWTFRH
IYRGQPKRHLLTTGWSVFVSTKRLFAGDSVLFVRDEKSQLMLGIRRANRQTPTLSSSVIS
SDSMHIGILAAAAHANANSSPFTIFFNPRASPSEFVVPLAKYNKALYAQVSLGMRFRMMF
ETEDCGVRRYMGTVTGISDLDPVRWKGSQWRNLQVGWDESTAGDRPSRVSIWEIEPVITP
FYICPPPFFRPKYPRQPGMPDDELDMENAFKRAMPWMGEDFGMKDAQSSMFPG**LSLVQWM**
**SMQQNNPLSGSATPQLPSALSSFNLPNNFASNDPSKLLNFQSPNLSSANSQFNKPNTVNH**
**ISQQMQAQPAMVKSQQQQQQQQQQHQHQQQQLQQQQQLQMSQQQVQQQGIYNNGTIAVAN**
**QVSCQSPNQPTGFSQSQLQQQSMLPTGAKMTHQNINSMGNKGLSQMTSFAQEMQFQQQLE**
**MHNSSQLLRNQQEQSSLHSLQQNLSQNPQQLQMQQQSSKPSPSQQLQLQLLQKLQQQQQQ**
**QSIPPVSSSLQPQLSALQQTQSHQLQQLLSSQNQQPLAHGNNSFPASTFMQPPQIQVSPQ**
**QQGQMSNKNLVAAGRSHSGHTDGEAPSCSTSPSANNTGHDNVSPTNFLSRNQQQGQ**AASV
SASDSVFERASNPVQELYTKTESRISQGMMNMKSAGEHFRFKSAVTDQIDVSTAGTTYCP
DVVGPVQQQQTFPLPSFGFDGDCQSHHPRNNLAFPGNLEAVTSDPLYSQKDFQNLVPNYG
NTPRDIETELSSAAISSQSFGIPSIPFKPGCSNEVGGINDSGIMNGGGLWPNQTQRMRTY
TKVQKRGSVGRSIDVTRYSGYDELRHDLARMFGIEGQLEDPLTSDWKLVYTDHENDILLV
GDDPWEEFVNCVQNIKILSSVEVQQMSLDGDLAAIPTTNQACSETDSGNAWKVHYEDTSA
AASFNR

**>ARF19 QtoS**

MKAPSNGFLPSSNEGEKKPINSQLWHACAGPLVSLPPVGSLVVYFPQGHSEQVAASMQKQ
TDFIPNYPNLPSKLICLLHSVTLHADTETDEVYAQMTLQPVNKYDREALLASDMGLKLNR
QPTEFFCKTLTASDTSTHGGFSVPRRAAEKIFPPLDFSMQPPAQEIVAKDLHDTTWTFRH
IYRGQPKRHLLTTGWSVFVSTKRLFAGDSVLFVRDEKSQLMLGIRRANRQTPTLSSSVIS
SDSMHIGILAAAAHANANSSPFTIFFNPRASPSEFVVPLAKYNKALYAQVSLGMRFRMMF
ETEDCGVRRYMGTVTGISDLDPVRWKGSQWRNLQVGWDESTAGDRPSRVSIWEIEPVITP
FYICPPPFFRPKYPRQPGMPDDELDMENAFKRAMPWMGEDFGMKDAQSSMFPGLSLVSWM
SMSSNNPLSGSATPSLPSALSSFNLPNNFASNDPSKLLNFSSPNLSSANSSFNKPNTVNH
ISSSMSASPAMVKSSSSSSSSSSSHSHSSSSLSSSSSLSMSSSSVSSSGIYNNGTIAVAN
SVSCSSPNSPTGFSSSSLSSSSMLPTGAKMTHSNINSMGNKGLSSMTSFASEMSFSSSLE
MHNSSSLLRNSSESSSLHSLSSNLSSNPSSLSMSSSSSKPSPSSSLSLSLLSKLSSSSSS
SSIPPVSSSLSPSLSALSSTSSHSLSSLLSSSNSSPLAHGNNSFPASTFMSPPSISVSPS
SSGSMSNKNLVAAGRSHSGHTDGEAPSCSTSPSANNTGHDNVSPTNFLSRNSSSGSAASV
SASDSVFERASNPVSELYTKTESRISSGMMNMKSAGEHFRFKSAVTDSIDVSTAGTTYCP
DVVGPVSSSSTFPLPSFGFDGDCSSHHPRNNLAFPGNLEAVTSDPLYSSKDFSNLVPNYG
NTPRDIETELSSAAISSQSFGIPSIPFKPGCSNEVGGINDSGIMNGGGLWPNQTQRMRTY
TKVQKRGSVGRSIDVTRYSGYDELRHDLARMFGIEGQLEDPLTSDWKLVYTDHENDILLV
GDDPWEEFVNCVQNIKILSSVEVQQMSLDGDLAAIPTTNQACSETDSGNAWKVHYEDTSA
AASFNR

**>ARF19 QtoG**

MKAPSNGFLPSSNEGEKKPINSQLWHACAGPLVSLPPVGSLVVYFPQGHSEQVAASMQKQ
TDFIPNYPNLPSKLICLLHSVTLHADTETDEVYAQMTLQPVNKYDREALLASDMGLKLNR
QPTEFFCKTLTASDTSTHGGFSVPRRAAEKIFPPLDFSMQPPAQEIVAKDLHDTTWTFRH
IYRGQPKRHLLTTGWSVFVSTKRLFAGDSVLFVRDEKSQLMLGIRRANRQTPTLSSSVIS
SDSMHIGILAAAAHANANSSPFTIFFNPRASPSEFVVPLAKYNKALYAQVSLGMRFRMMF
ETEDCGVRRYMGTVTGISDLDPVRWKGSQWRNLQVGWDESTAGDRPSRVSIWEIEPVITP
FYICPPPFFRPKYPRQPGMPDDELDMENAFKRAMPWMGEDFGMKDAQSSMFPGLSLVGWM
SMGGNNPLSGSATPGLPSALSSFNLPNNFASNDPSKLLNFGSPNLSSANSGFNKPNTVNH
ISGGMGAGPAMVKSGGGGGGGGGGHGHGGGGLGGGGGLGMSGGGVGGGGIYNNGTIAVAN
GVSCGSPNGPTGFSGSGLGGGSMLPTGAKMTHGNINSMGNKGLSGMTSFAGEMGFGGGLE
MHNSSGLLRNGGEGSSLHSLGGNLSGNPGGLGMGGGSSKPSPSGGLGLGLLGKLGGGGGG
GSIPPVSSSLGPGLSALGGTGSHGLGGLLSSGNGGPLAHGNNSFPASTFMGPPGIGVSPG
GGGGMSNKNLVAAGRSHSGHTDGEAPSCSTSPSANNTGHDNVSPTNFLSRNGGGGGAASV
SASDSVFERASNPVGELYTKTESRISGGMMNMKSAGEHFRFKSAVTDGIDVSTAGTTYCP
DVVGPVGGGGTFPLPSFGFDGDCGSHHPRNNLAFPGNLEAVTSDPLYSGKDFGNLVPNYG
NTPRDIETELSSAAISSQSFGIPSIPFKPGCSNEVGGINDSGIMNGGGLWPNQTQRMRTY
TKVQKRGSVGRSIDVTRYSGYDELRHDLARMFGIEGQLEDPLTSDWKLVYTDHENDILLV
GDDPWEEFVNCVQNIKILSSVEVQQMSLDGDLAAIPTTNQACSETDSGNAWKVHYEDTSA
AASFNR

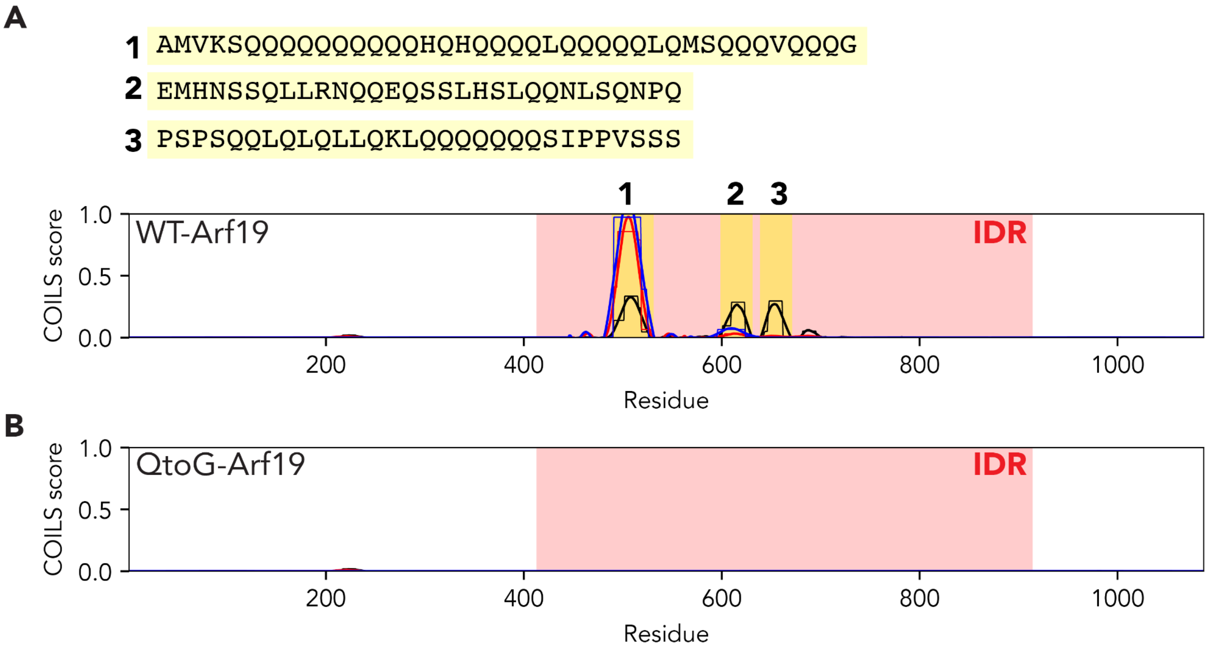


**Figure S1. QtoG mutant ablates any possible coiled coils. (A)** Coiled-coil prediction using the COILS web server (https://embnet.vital-it.ch/software/COILS_form.html) identifies three possible regions that may contain predicted coiled-coils, highlighted as regions 1, 2, and 3 with the sequences shown above. Different color lines represent different sized sliding windows (black = 14, red = 21, blue = 28). Coiled-coil prediction analysis was performed using server default values. The Arf19 IDR is highlighted in red. **(B)** Identical analysis performed on the QtoG variant reveals loss of predicted coiled coils across the IDR.

| | % Monomer | % Dimer | % Trimer | % Tetramer |
|---|---|---|---|---|
| Average | 75.403 | 20.842 | 2.959 | 0.796 |
| Standard Deviation | 24.542 | 19.842 | 3.830 | 1.111 |

**Table S1. The ARF19 PB1 domain alone does not form higher-order oligomers in protoplasts.** This table contains average values and the standard deviation for various oligomers quantified using Number and Brightness analysis of protoplasts expressing the ARF19 Phox/Bemp1 domain tagged with mVenus. N = 7.
